# Photoprotective Bioactive Pigment Produced by *Bacillus altitudinis* MIM2 Isolated from Mundra Port, Kutch, Gujarat, India

**DOI:** 10.1101/2022.03.15.484403

**Authors:** Pankhaniya Mira Gordhanbhai, Chavda Jigna, Murshidah M Shereen, Noble K Kurian

## Abstract

Melanins are ubiquitous black or brown color pigments exhibiting a wide variety of bioactivities. They are stable and insoluble in nature. Melanin is an industrially important pigment currently used in the cosmetics, medicine and pharmaceutical industries. Bacteria secrete melanin extracellularly, which makes the downstream processing of bacterial melanin easier. This makes bacteria a good source of melanin. Stress conditions like salt stress, radiation stress, etc., trigger the bacteria to produce melanin and several bacterial species were found to produce melanin under different stress-induced conditions. In this present study, we isolated melanin-producing bacteria from a saline sediment sample of Mundra port, Kutch region of Gujarat state of India. The melanin-producing bacteria was characterized by staining and biochemical and molecular methods. The melanin produced was extracted and analyzed using physicochemical techniques. The extracted melanin had shown good anti-bacterial and radical scavenging activity. This melanin is also shown to protect the bacteria from UV irradiation. To our knowledge, this is the first report on *Bacillus altitudinis* producing melanin.

## INTRODUCTION

Melanins are bioactive pigments produced by most living organisms. Its function mainly ranges from photoprotection to contributing to virulence in these organisms^1^. These black pigments can be classified mainly into three types based on their difference in biosynthetic process. They are eumelanin, synthesized via DOPA, while cysteine is incorporated into DOPA in the second type of melanin, i.e., pheomelanin. All other remaining types of melanin are classified under allomelanins synthesized from a wide variety of substrates. Allomelanins include bacterial pyomelanin, fungal DHN melanins, plant catechol melanin and so on ^2^. Though the melanins originate via different metabolic pathways, they exhibit similar physicochemical properties. The properties include their nonpolar nature, effective antioxidant activity, efficient metal chelating, and radioprotective activity, which make them a good candidate for many useful applications.

Extraction of melanin is a complex process in many organisms as contamination of cellular materials could occur in the pigment, especially protein contamination. Bacteria produce melanin extracellularly in culture medium, making it easier for the downstream processing of the pigment than melanin from other organisms. This makes bacterial melanin preferred over melanin from other sources. The major drawback of bacterial melanin is that bacteria only produce a low quantity of pigment. So, the search for better melanin-producing bacteria is still a major research topic in bioprocess technology ^3^.

Melanin has been discovered to neutralize the oxidants produced by environmental stress. Melanized *Cryptococcus neoformans* survived in nitrogen and oxygen-derived reactive intermediates 10 times better than non-melanized cells. Similar protection from reactive oxygen species was reported in *Wangiella dermatitis* and *Alternaria alternata*. Melanin was found to protect melanocytes and keratinocytes from hydrogen peroxide-mediated DNA damage. This reflects the effective protective property of melanin against free radicals. UV light can induce the production of reactive oxygen species, which will ultimately lead to tissue damage or induction of diseases like skin cancer. Besides being an antioxidant, melanin can confer UV protective properties, making it an ideal choice in cosmetic formulations ^4^.

Stress usually induces bacteria to produce melanin. Bacteria isolated from stress-prone areas are considered to be good candidate strains to be screened for melanin production. In the present research, melanin-producing bacteria were isolated from the saline soil of the Kutch region, and the melanin produced was characterized. The antioxidant, antimicrobial, and UV protective properties of the pigment were explored further.

## MATERIALS & METHODS

### Isolation of bacteria from saline soil

The bacteria were isolated from the saline soil of Mundra Port, Kutch Region (22.74°N, 69.7°E), Gujarat, India. Saline soil was collected in autoclaved polythene bags and brought to the laboratory. One gram of the soil was serially diluted, and dilutions were spread and plated on nutrient agar containing 4% sodium chloride to select halophilic bacteria. The bacteria grown in the plates were purified using quadrant streaking and used for screening for further melanin production.

### Screening for melanin production

In boiling tubes, the isolated bacteria were screened for melanin production in the tyrosine basal broth ^5^ (5 mL). The bacteria that produced a good amount of melanin are further inoculated in 100 mL tyrosine basal broth for production. 1 OD 5% culture was inoculated, and media was incubated at 37°C for 8-10 days. After 10 days of incubation, the tyrosine basal broth was centrifuged, and the bacterial pellet was removed. The supernatant was then read spectrophotometrically at 400 nm ^6^, and the concentration of melanin produced was determined using a standard graph made of synthetic melanin.

### Characterization of melanin-producing bacteria

Melanin-producing bacteria were characterized using Gram staining and biochemical tests ^7^. Further, the bacteria were tested for antibiotic susceptibility using the Kirby-Bauer disc diffusion method ^8^. The MAR (Multiple Antibiotic Resistance) index was calculated as the ratio between the number of antibiotics an isolate is resistant to and the total number of antibiotics tested ^9^. The bacteria were identified using 16S rDNA sequencing at NCIM, Pune, India. The obtained sequence was then searched using the BLAST tool at NCBI to identify the bacteria. Subsequently, the sequence was submitted to GenBank, and an accession number was obtained. The phylogenetic tree of the melanin-producing bacteria and related species was constructed using the MEGA 7 software and the Neighbour-joining method ^10^.

### Extraction and purification of melanin

After 10 days of incubation, the production medium is centrifuged (BioEra, Japan) at 6000 rpm for 10 minutes at 28°C to separate the bacterial pellet from the supernatant containing melanin. The supernatant is then further acidified to below pH 2 with 1N HCl. As the pH decreases, a black melanin precipitate is seen at the bottom of the flask. The mixture is then centrifuged again at 6000 rpm for 10 minutes at 37°C to pellet the melanin. The resulting pellet is washed twice with absolute ethanol and distilled water to obtain purified melanin^11^. The purified melanin is then dried in a hot air oven (Equitron oven - steam series, India) for 2 days at 80°C.

### Characterization of melanin

#### Solubility of melanin in solvents

The solubility of melanin in water, ethanol, methanol, isopropanol, acetic acid, HCl, H_2_SO_4_, DMSO, and NaOH was tested. Melanin powder was added and vortexed for 5 minutes, then centrifuged at 5000rpm for 1 minute. The results were observed to check whether the melanin dissolved ^12^.

#### UV-Visible Spectroscopy

UV-visible spectra were obtained by scanning the melanin solution from 190 nm to 890 nm using a UV-visible spectrophotometer (Jasco V-730 Spectrophotometer, Japan). The results were compared to synthetic melanin and earlier reports to confirm the pigment’s identity as melanin ^13^.

#### Fourier-Transform Infrared (FT-IR) Spectroscopy

The FT-IR spectrum is recorded at 4,000-400 cm^-1^ ^14^. Characteristic peaks obtained were compared with earlier reports.

#### Particle size and zeta potential analysis

The melanin is dispersed in Milli-Q water and sonicated for 10 min before the measurement. Dispersed melanin samples were observed in the concentration of 100µL/1mL. Size distribution and surface charge of the pigment was determined by the dynamic light scattering (DLS) and zeta potential instrument (Zetasizer Nano ZS, Malvern) at 25 °C

#### Antioxidant activity of melanin

The antioxidant or free-radical scavenging activity of melanin was determined using the 2,2-Diphenyl-1-picrylhydrazyl (DPPH) free radical scavenging assay as per Liyana and Shahidi, 2005 ^15^. Ascorbic acid served as the positive control. Different concentrations of melanin were added to DPPH and incubated in the dark for 30 minutes. The absorbance of the test samples was measured spectrophotometrically at 517 nm.

The ability to scavenge DPPH radicals was calculated by the equation

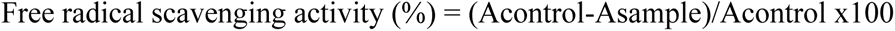

Acontrol is the absorbance of the DPPH + methanol, and Asample is absorbance of the free radical solution with melanin/standard antioxidant.

#### Antibacterial activity of melanin

The antibacterial activity of melanin was tested against the bacteria *Bacillus* sp., *E. coli, Salmonella* sp., *Shigella* sp., and *Staphylococcus aureus* using a disc diffusion assay. The bacteria were swabbed on Mueller-Hinton agar plates, and melanin at different concentrations (10μg/mL, 20μg/mL, 40μg/mL, 60μg/mL, 80μg/mL, 100μg/mL) was poured into wells bored in the plates. Streptomycin was used as the control in the study. The plates were then incubated at 37°C for 24 hours, and clear zone formation around the wells was checked ^16^.

#### UV photoprotection activity of melanin

Melanin-producing and non-melanin-producing MIM2 strains were exposed to shortwave UV radiation of 254 nm from a laminar airflow UV lamp. The melanin (tyrosine basal broth) and non-melanin (nutrient broth) bacteria were positioned 30 cm away from the UV lamp of the laminar airflow. The UV treatment was administered for the following durations: 0 minutes, 2 minutes, 5 minutes, 10 minutes, and 20 minutes. After treatment, 100μl of the broth was spread on nutrient agar plates and incubated for 24 hours at 37°C. The following day, both samples were assessed for differences in bacterial growth ^17^.

#### Sun Protection Factor (SPF) analysis

Sun protection factor of melanin was determined as per Mansur equation^31^. 100µg/mL melanin in 1mL ethanol is used for the study. The blank was 100% ethanol for this analysis^32^.

## RESULT & DISCUSSION

### Isolation of melanin-producing bacteria

A total of 30 colonies were obtained in the pour-plated plates, which were subjected to quadrant streaking to obtain pure colonies (Figure 1(a)). In test tubes, the pure colonies were then tested for melanin production in 5 mL of tyrosine basal broth. After 10 days of incubation, one tube (MIM2) showed a significant color change from white to dark brown (Figure

**Figure 1.**
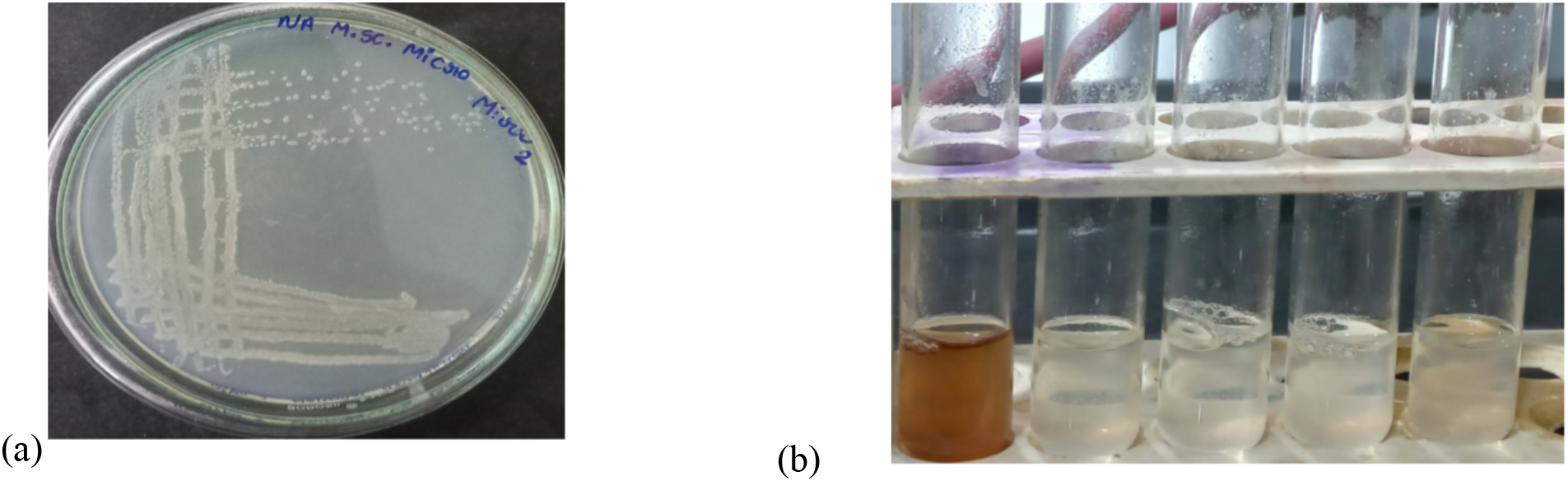
(a) Nutrient agar plates showing MIM2 colonies (b) tyrosine basal broth tube with MIM2 showing melanin production

### Identification and Characterization of melanin-producing bacteria

The bacteria MIM2 was characterized as Gram-positive rods (Figure 2). The Indole, Voges-Proskauer, and Citrate tests were found to be negative, while the Methyl Red and catalase tests were positive (Table 1). Genotypic characterization was performed using 16S rDNA sequencing, and the sequence analysis was conducted using NCBI Nucleotide BLAST tool ^18^. The organism MIM2 has been identified as *Bacillus altitudinis*, and the sequence has been submitted to GenBank. The accession number (OM967420) has been obtained. A phylogenetic tree (Figure 3) was constructed using the neighbour-joining method ^19^ and the nucleotide-based TN84 evolutionary model for estimating genetic distances based on synonymous and nonsynonymous nucleotide substitutions using the software MEGA 7. Statistical support for branching was estimated using 1000 bootstrap steps. The phylogenetic tree reveals *that Bacillus altitudinis* strain MIM2 is similar to salt-tolerant bacteria like *Bacillus atrophaeus.* This confirms that strain MIM2 is halophilic in nature.

**Figure 2:**
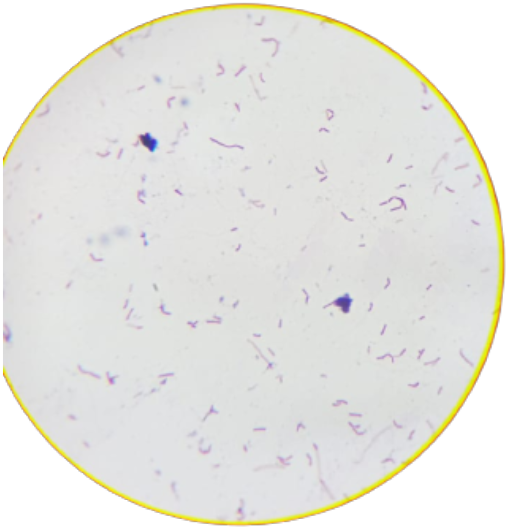
Gram staining image of MIM2 (40X) showing Gram-positive rods

**Figure 3:**
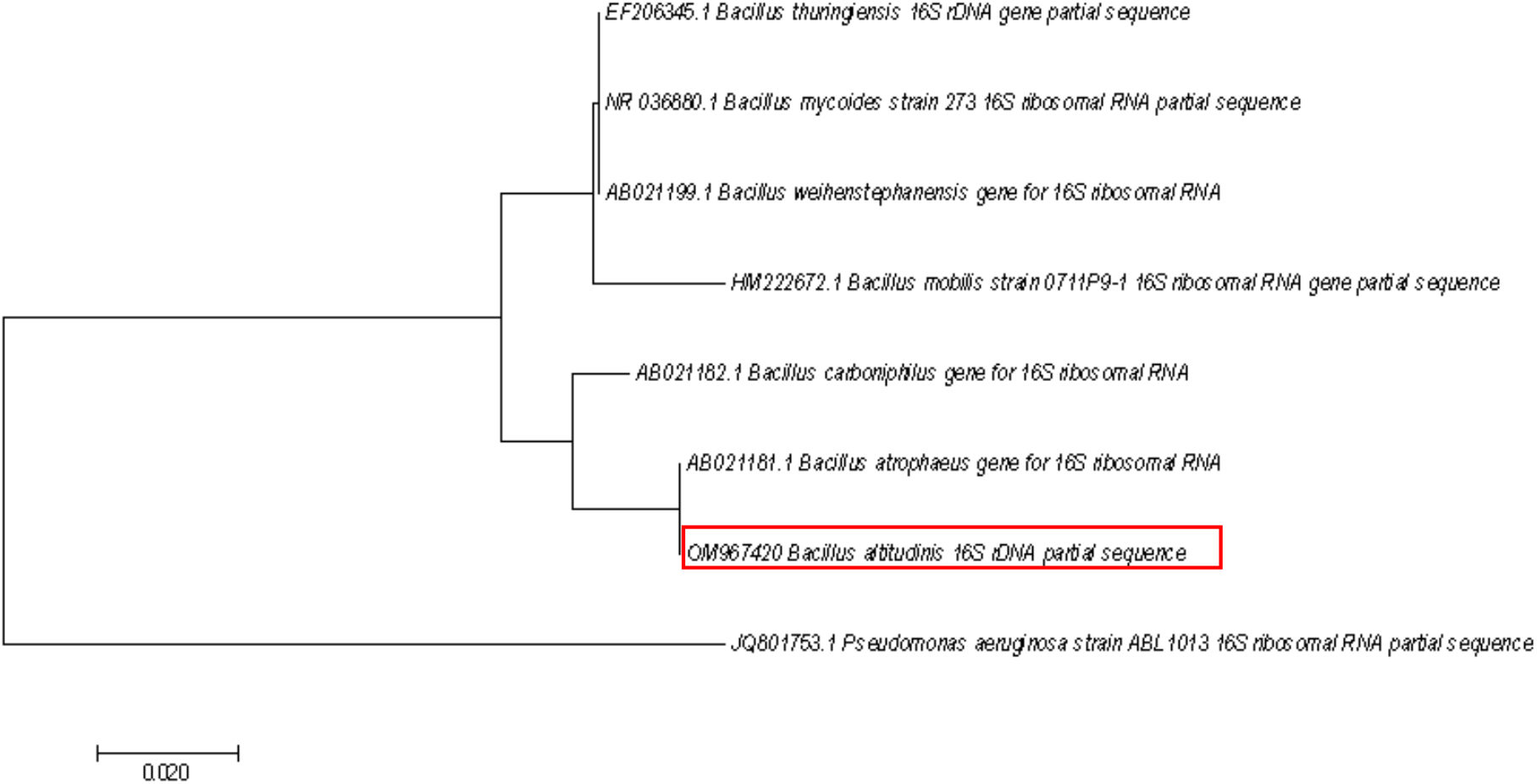
The phylogenetic tree of MIM2 inferred using the Neighbor-Joining method.

**Table 1:** Biochemical Characterization of MIM2.

| Biochemical test | Results |
| --- | --- |
| Indole | - |
| Methyl Red | + |
| Voges-Proskauer | - |
| Simmons Citrate | - |
| Catalase | + |

Strain MIM2 had shown sensitivity to the majority of the antibiotics tested. The strain had shown resistant to CF: Cefotaxime and Cl: Ceftizoxime only (Figure 4, Table 2). The MAR index was found to be 0.16. this indicates that the sampling area was not contaminated with antibiotics.

**Figure 4:**
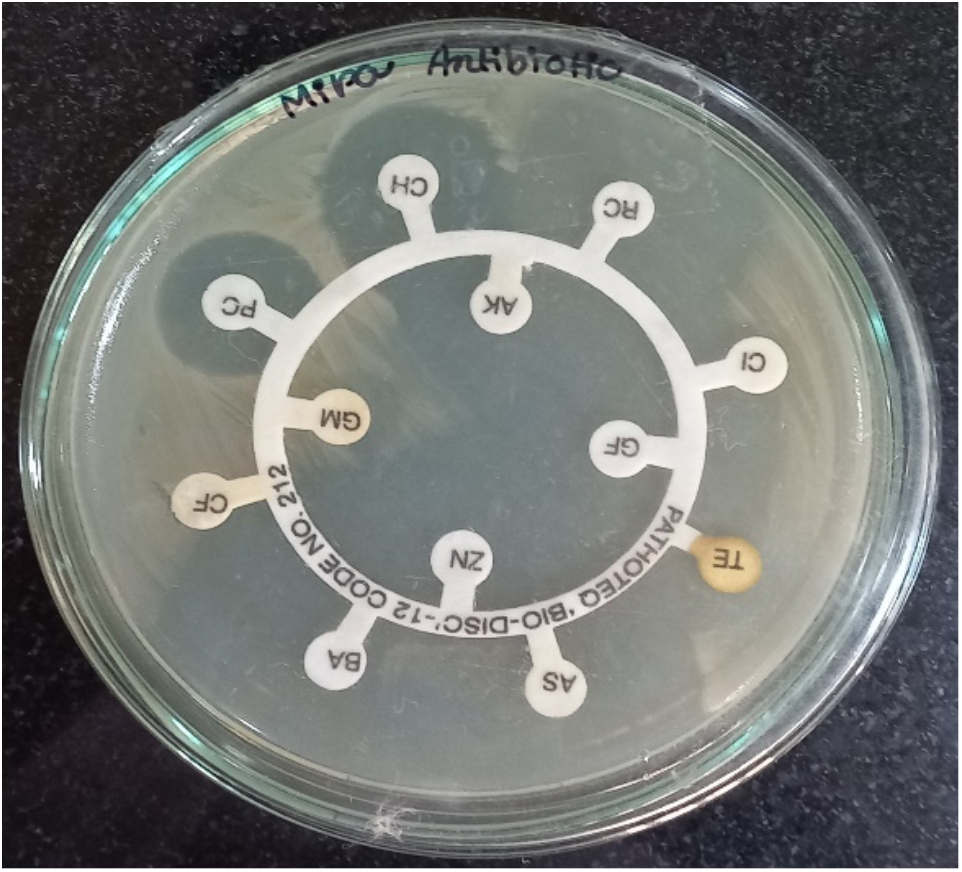
Antibiotic sensitive tests for the strain MIM2

**Table 2:** Antibiotic sensitivity profile of strain MIM2.

|  | Sensitive (+) |
| --- | --- |
| Antibiotics | Resistant (-) |
| AS: Ampicillin | + |
| BA: Co-Trimoxazole | + |
| CF: Cefotaxime | - |
| PC: Piperacillin | + |
| CH: Chloramphenicol | + |
| RC: Ciprofloxacin | + |
| Cl: Ceftizoxime | - |
| TE: Tetracycline | + |
| ZN: Ofloxacin | + |
| GM: Gentamicin | + |
| AK: Amikacin | + |
| GF: Gatifloxacin | + |

### Melanin production

To scale up the melanin production, 1 OD culture of MIM2 was inoculated in tyrosine basal broth, and melanin production was monitored every 12 hours. Color change was observed from the 6^th^ day of incubation, and it increased to 10 days of incubation. After the 10th day, the amount of melanin (Figure 5 (a)) produced was estimated spectrophotometrically at 400nm, and the concentration of melanin produced was quantified using a synthetic melanin standard curve. The amount of melanin produced by MIM2 was 245.67±8.7 mg/L. The amount of melanin produced is comparable with previous reports. *Bacillus* sp. is known to produce melanin in many instances. For example, *Bacillus* sp. BTCZ31 was reported to produce 32.63±0.4 μg/mL ^20^ of melanin, while *Bacillus subtilis* 4NP-BL produced 1.5 g dry wt L–1 of melanin ^21^. A strain of *Bacillus thuringiensis* BMB181 was found to produce 8.55 mg/mL of melanin ^22^. Compared to these reported *Bacillus* sp. strains, MIM2 produced a moderate amount of melanin without optimization of media components. It is likely that optimizing the media components could enhance the melanin production by the bacteria.

**Figure 5:**
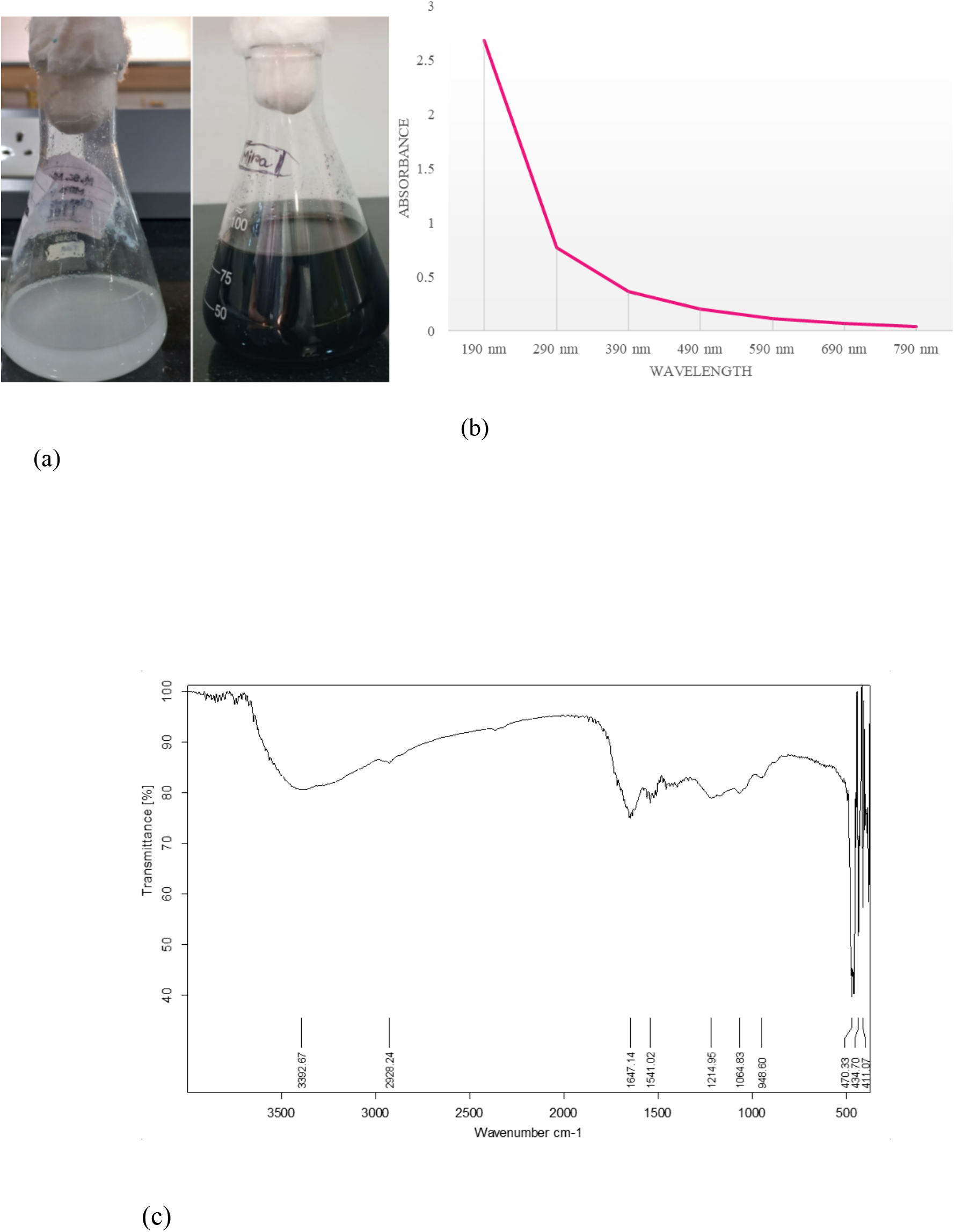
(a) Melanin production in tyrosine basal broth by MIM2 (b) UV Visible spectrum MIM2 melanin (c) FTIR spectrum of MIM2 melanin

### Characterization of Melanin

The chemical reactivity of MIM2 melanin was tested by testing its solubility in acidic, alkaline, and organic solvents (Table 3). Melanin was found to be soluble in 1 N NaOH only, and in DMSO, it was found to be sparingly soluble, with some particles remaining. The solubility of melanin in basic solutions is thought to arise from the deprotonation of the various acid/base groups present in melanin, including carboxylic acid and phenolic groups ^23^. Melanin insolubility is one of the factors that makes it different from other bacterial pigments such as carotenoids, etc.

**Table 3:** Solubility of melanin in various solvents.

| Solvent | Solubility |
| --- | --- |
| Distilled Water | Insoluble |
| 1N NaOH | Soluble |
| Ethanol | Insoluble |
| Methanol | Insoluble |
| DMSO | Sparingly Soluble |
| Isopropanol | Insoluble |
| Acetic Acid | Insoluble |
| **HCl** | Insoluble |
| **H<sub>2</sub>SO<sub>4</sub>** | Insoluble |

Purified melanin is analyzed using UV visible and FTIR spectrophotometry to confirm the pigment as melanin. The pigment shows a characteristic featureless UV-visible spectrum, with maximum absorption at the UV region that significantly decreases as it reaches the visible region (Figure 5 (b)). The spectrum was found similar to earlier reports ^24^.

FTIR spectrum of melanins (Figure 5 (c)) had shown considerable similarity with synthetic melanin and earlier reports ^25,26^. The spectrum showed a broad absorption around 3392 cm^-1^, corresponds to phenolic –OH and –NH stretching vibrations. Characteristic peaks observed between 1600-1400 cm^-1^ was attributed to aromatic ring C=C stretching. This confirmed the polyphenolic and aromatic nature of MIM2 melanin.

The halophilic bacteria *Bacillus altitudinis* MIM2 produces melanin pigment, as confirmed by spectroscopic characterization. Further characterization is required for potential application studies.

### Particle size and zeta potential analysis

The size of the MIM2 melanin particles was determined by dynamic light scattering. The hydrodynamic diameter of the melanin was found to be 377.3 nm [Figure 6]. The distribution curve spanned roughly 200 to 1000 nm. Particles in this size range was found similar to the earlier reports in melanin^33^. A particle size in the sub-micron range is relevant for cosmetic use. It could help the pigment disperse in lotions and creams and give uniform coverage on the skin surface, and it may be adjusted further by homogenization or sonication.

**Figure 6.**
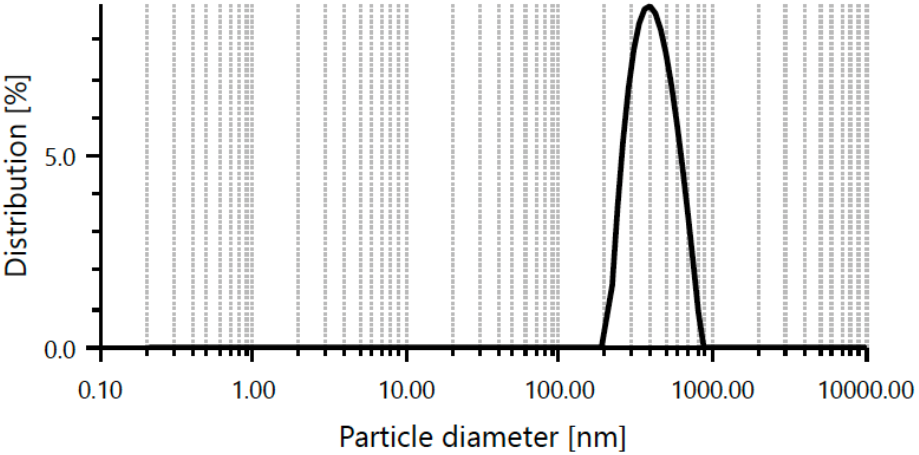
Particle size distribution (intensity) of MIM2 melanin determined by dynamic light scattering.

The surface charge of the melanin particles was assessed by zeta potential analysis [Figure 7]. The mean zeta potential was −16.1 mV. The negative charge is consistent with the ionizable carboxyl and phenolic hydroxyl groups of melanin, which carry a negative charge in aqueous medium. These groups were also indicated by the FT-IR spectrum (broad peak at 3500 cm⁻¹ area for phenolic –OH/–NH). The magnitude of the zeta potential is, however, low. Colloids with zeta potential values beyond ±30 mV are generally considered electrostatically stable, while values close to zero indicate weak repulsion between particles and a tendency to aggregate or sediment over time. The low zeta potential of MIM2 melanin therefore suggests that the particles are only weakly stabilized by electrostatic repulsion. This agrees with the water-insoluble nature of the pigment (Table 3).

**Figure 7.**
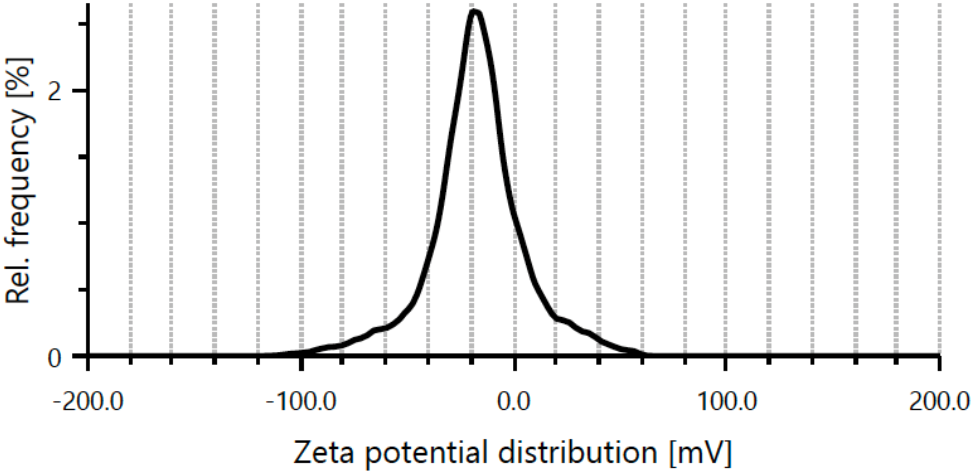
Zeta potential distribution of MIM2 melanin.

### Antioxidant activity of melanin

The free radical scavenging activity of MIM2 melanin was assayed using DPPH method ^15^. MIM2 melanin was shown to have very good radical scavenging activity of 66.4±1.1% (100 µg/mL) compared to control ascorbic acid, which was 92.16±1.8% for the same concentration (Figure 8). A thermophilic *Bacillus* sp was found to show 100% antioxidant activity earlier ^27^. Another endophytic *Bacillus subtilis* had shown 94.47% at 100 μg mL concentration ^28^. Though MIM2 melanin showed a moderate amount of antioxidant activity compared to the control, it was found to be significant.

**Figure 8:**
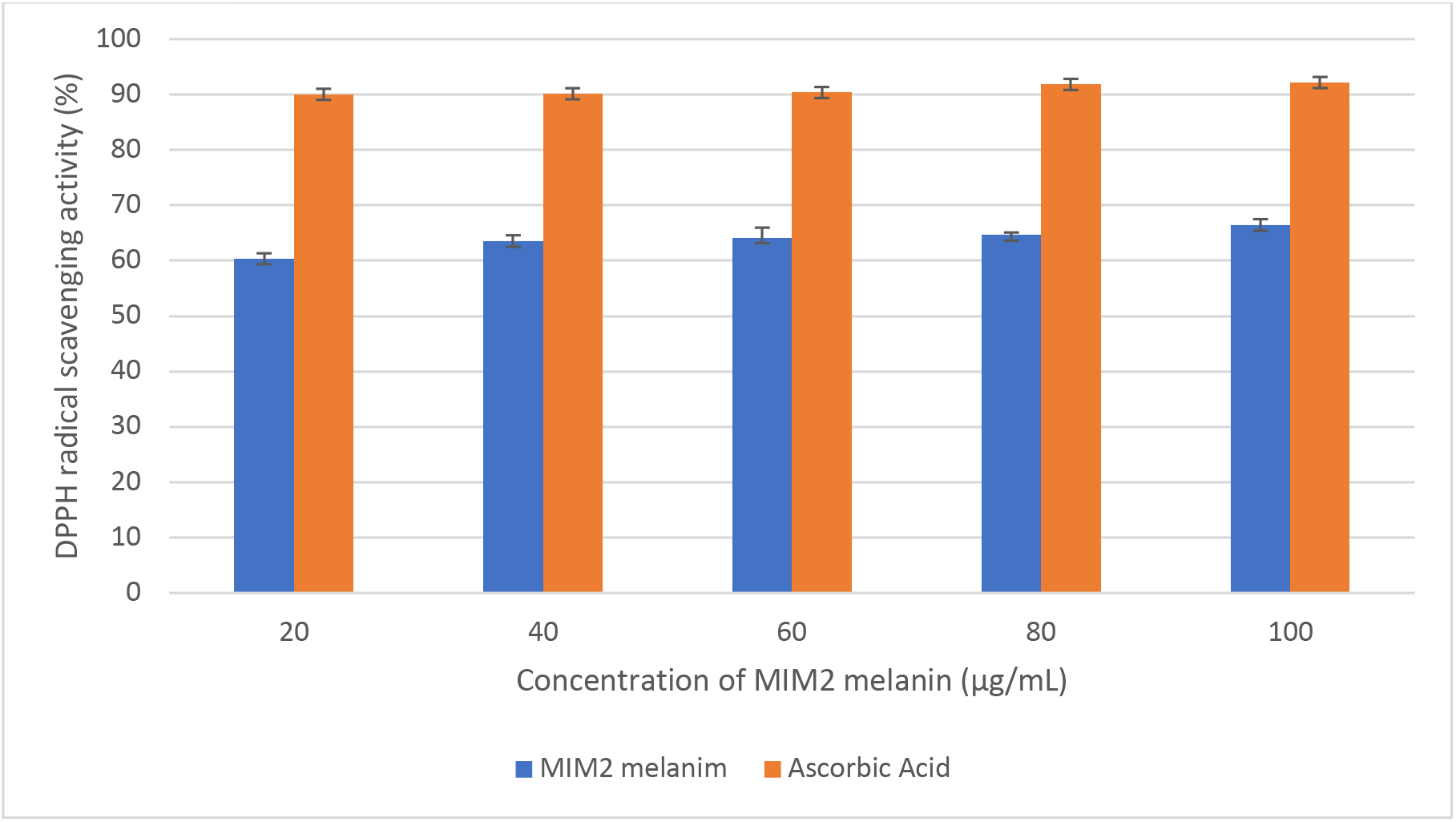
Radical scavenging activity of MIM2 melanin

### Antibacterial activity of melanin

The antibacterial activity of the melanin was evaluated using well diffusion assay. MIM2 melanin had shown good antibacterial activity at even lower concentrations against potential pathogens. The minimum inhibitory concentration was between the range of 20-40μg/mL against all bacteria tested (Table 4). <u>Lachnum</u> YM30 intracellular melanin has shown antibacterial activity against Gram-negative bacteria *Escherichia coli, Salmonella typhi, Vibrio parahaemolyticus* and Gram-positive bacteria *Listeria monocytogenes, Bacillus megaterium,, Staphylococcus aureus* ^29^ at similar concentration to that of the current study. This indicates that melanin could be an effective antibacterial agent against potential pathogens.

**Table 4:**
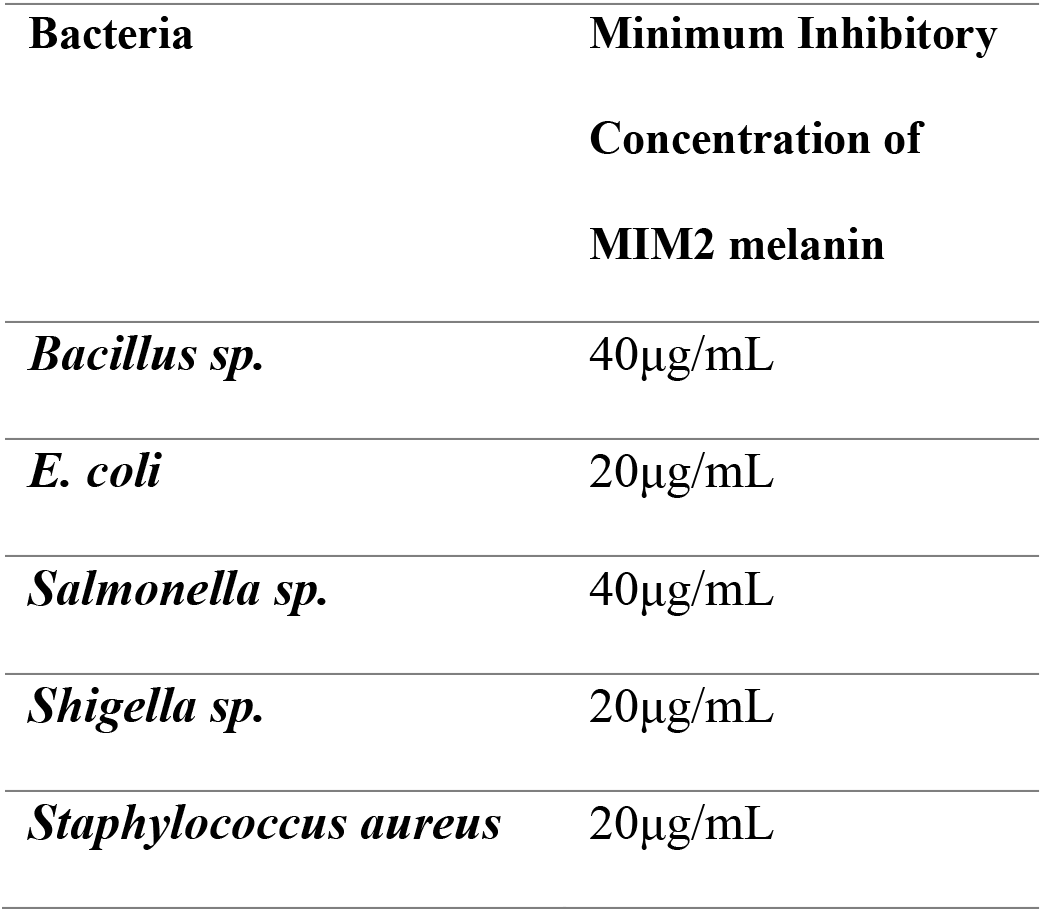
Antibacterial activity of MIM2 melanin.

### UV photoprotection of melanin

After 24 hours of incubation, the melanized colony plate showed more colonies than the non-melanized strain MIM2. The number of colonies was too high to count on both plates, but the difference was evident visually. There was a significant difference in the counts after 20 minutes of UV treatment on a non-melanized bacterial plate (Figure 7b). In contrast, melanized bacterial plates contained colonies beyond the countable limit, mostly seen in patches that cannot be differentiated into individual colonies (Figure 7a). This variation was observed in all plates for all durations of UV treatment. Qualitatively, the change was evaluated from the Petri plates (Figure 9), indicating that melanized colonies are more protected from UV radiation, as denoted by the increased colony count.

**Figure 9:**
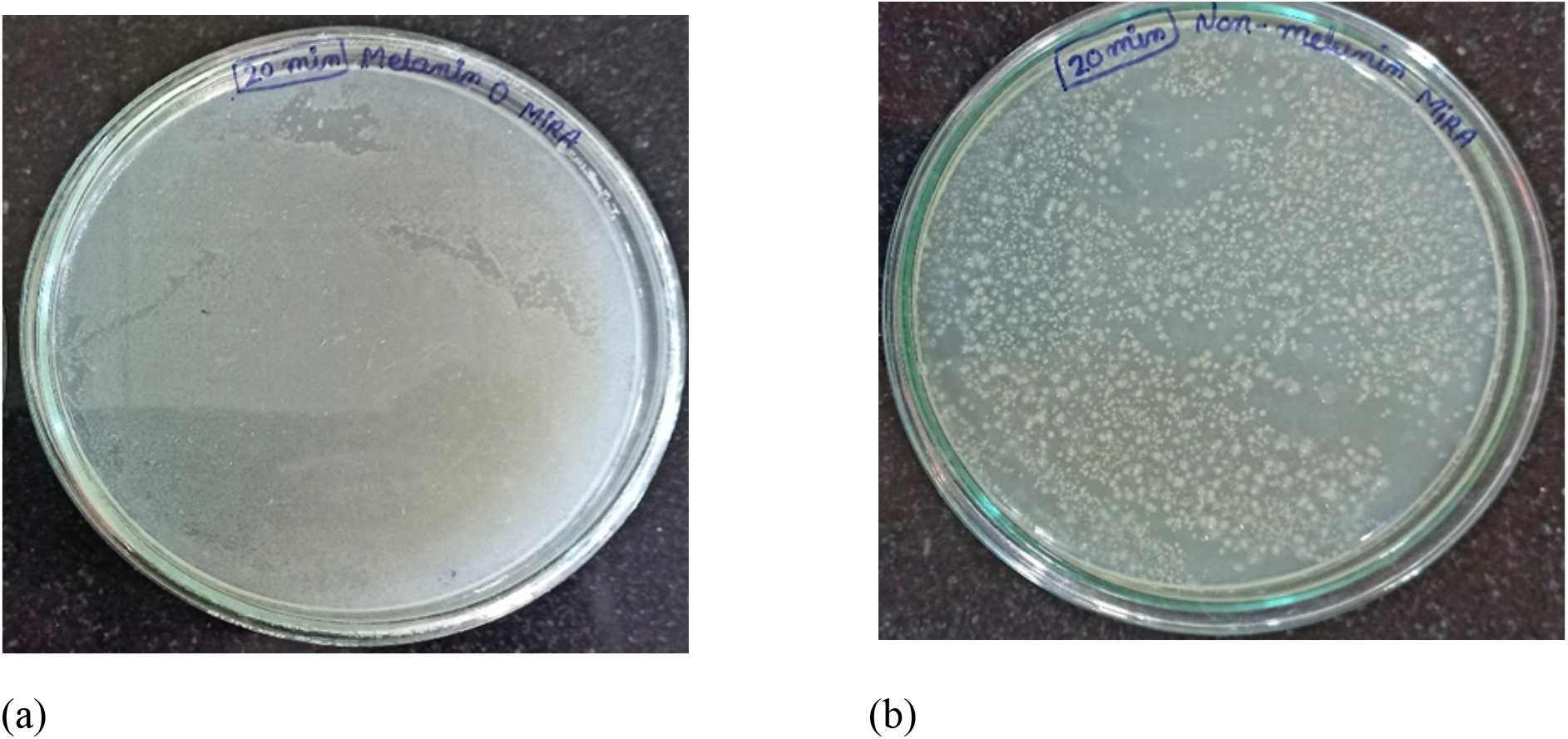
UV protection of (a) melanin-producing and (b) non-melanin-producing bacterial growth on Nutrient agar media

According to Joshi et al., 2021 ^17^, after up to 15 minutes of UV treatment, melanized cells resisted and survived. However, in the present study, the melanized MIM2 strain survived even after 20 minutes of UV treatment. Melanin has been shown to enhance the SPF value of sunscreen cosmetics in many earlier reports ^30^. Photoprotection is an important property for most cosmetic products; MIM2 melanin, with its immense potential, could be a good ingredient in cosmetic solutions.

### Sun Protection Factor (SPF) analysis

The SPF value of MIM2 melanin by an average of 14.17 ±1.6 (Table 5). SPF values fall in the range of moderate sun protection (SPF 12–<30) in the commonly used classification. Considering that the melanin was used without any other UV filter, this supports the photoprotective potential of MIM2 melanin and agrees with the colony survival after UV treatment in the present study (Figure 9).

**Table 5:** Sun Protection Factor Calculation.

| Wavelength | EE($\lambda$)×I($\lambda$) | F1 | | F2 | |
| --- | --- | --- | --- | --- | --- |
| | | Abs | Abs × EE($\lambda$)× I($\lambda$) | Abs | Abs × EE($\lambda$)× I($\lambda$) |
| 290 | 0.015 | 1.645 | 0.024675 | 1.403 | 0.021045 |
| 295 | 0.0817 | 1.602 | 0.1308834 | 1.366 | 0.1116022 |
| 300 | 0.2874 | 1.564 | 0.4494936 | 1.334 | 0.3833916 |
| 305 | 0.3278 | 1.529 | 0.5012062 | 1.305 | 0.427779 |
| 310 | 0.1864 | 1.485 | 0.276804 | 1.273 | 0.2372872 |
| 315 | 0.0837 | 1.440 | 0.120528 | 1.243 | 0.1040391 |
| 320 | 0.018 | 1.394 | 0.025092 | 1.194 | 0.021492 |
|  | Total |  | 1.5286822 |  | 1.3066361 |
|  | SPF |  | 15.286822 |  | 13.066361 |

## CONCLUSION

The black pigment melanin from *Bacillus altitudinis* MIM2 has been found to have profound bioactivities, which could be utilized in many applications. Its immense antioxidant activity will help in use in cosmetic formulations as a tissue protectant. Its photoprotective nature will help protect from UV-mediated injury and further chances of getting skin cancer-like dreadful diseases. The antibacterial activity of the pigments adds to the formulations in preventing pathogenic bacteria and helps act as a preservative in its storage. Though melanin was reported as non-cytotoxic, MIM2 melanin must be evaluated for its cytotoxicity before being utilized in different applications.

